# Coherent Structured Illumination Microscopy with Enhanced Optical Sectioning

**DOI:** 10.64898/2026.06.11.731428

**Authors:** Kevin T. Crampton, Alan G. Joly, Long D. Nguyen, Saleem Iqbal, Robert W. Boyd, James E. Evans

## Abstract

Coherent structured illumination microscopy (c-SIM) is a synthetic aperture optical technique for sub-diffraction limit imaging that extends the utility of traditional SIM to non-fluorescent samples. Here, we present a complementary 5-beam implementation of c-SIM that provides enhanced optical sectioning compared to conventional quadrupolar illumination. Since our approach detects intensity images due to coherent light scattering, it avoids the complications associated with detecting complex fields. Through comparative measurements on calibration samples and live microalgae, we show that 5-beam c-SIM effectively suppresses coherent defocus effects, improving image quality while simultaneously providing a 2-fold lateral resolution improvement.

## 1. Introduction

Structured illumination microscopy (SIM) is a well-established technique for improving the spatial resolution of wide-field (WF) microscopes and is most commonly applied to fluorescence imaging.[1] In its most basic implementation, SIM involves using programmable electro-optical devices or gratings to synthesize periodic illumination patterns out of coherent or partially coherent light. Periodic illumination shifts specimen frequencies that would normally fall outside the optical frequency support of an imaging system into the detection window. These higher spatial frequencies can be unmixed from the canonical sample frequencies and reconstructed into a final image with up to a factor-of-2 resolution improvement beyond the diffraction limit for fluorescence.[2-5] 3D volumetric imaging implementations have been demonstrated as well as non-linear versions that further improve the attainable spatial resolution by generating harmonics of the periodic illumination using saturated fluorophores.[6, 7]

While fluorescence-based SIM is widely used, supported on several commercial microscopy platforms, and complemented by numerous open-source reconstruction algorithms,[8-10] its application generally remains confined to fluorescently labeled or intrinsically fluorescent samples. In the more universal case of image formation through coherent light scattering, fluorescence-based SIM tools are no longer applicable because of the nonlinear relationship between the object field and the image intensity. Coherent structured illumination microscopy (c-SIM) has been developed to extend the utility of conventional SIM to scattering samples under coherent illumination.[11-15] A detailed theoretical framework for c-SIM as well as an experimental demonstration of 2-fold lateral resolution improvement in WF scattering imaging through structured oblique illumination microscopy (SOIM) was given by Chowdhury et al.[16] In contrast to conventional SIM, which typically involves illumination with re-oriented 1D sinusoidal patterns, SOIM relies on quadrupolar illumination to generate the 2D patterns required to completely capture the enhanced-resolution information. Methods for obtaining isotropic lateral resolution have been proposed[17] and demonstrated using high numerical aperture (NA) objective lenses.[18]

A key factor limiting the widespread adoption of c-SIM is its relatively poor optical sectioning (OS) ability. Compared to incoherent signal detection, whereby different emissions from the sample add but do not interfere, coherent signals contain interference cross-terms since the phases of the light scattered from various regions of the sample are fixed, not stochastic. Consequently, c-SIM provides up to Abbe diffraction-limited resolution in the lateral dimension, but not super-resolution as is the case for fluorescence-based SIM. This also influences the range of spatial frequencies accessible in the axial direction. Whereas the optical transfer function (OTF) in fluorescence imaging has finite axial extent, the coherent transfer function (CTF) is a thin locus in 3D Fourier space. As such, coherent-field contributions from different specimen depths are superposed without suppression, which leads to artifacts such as ringing around image features, speckle-like or grainy backgrounds, and aliasing. These effects are especially problematic for thick biological specimens such as cells or tissues. This has led to the development of several synthetic aperture techniques for imaging coherent reflectance or transmittance including optical diffraction tomography (ODT) and structured illumination-enabled 3D quantitative phase microscopy (SI-QPM).[19-25] Through different implementations, ODT and SI-QPM provide 3D refractive index or quantitative phase information by mapping out Fourier space with modulated illumination wavevectors. In ODT, this is accomplished by tilting the illuminating beam and recording images at many angles. Similarly, SI-QPM involves varying the structured illumination pattern frequency and acquiring multiple phase-stepped images for each carrier. These modalities typically rely on interferometric detection to recover the complex image field. The overall 3D spatial-frequency support is extended in both techniques since ODT combines measurements at multiple oblique illumination angles (tomographic filling) and SI-QPM heterodynes sample frequencies previously inaccessible to the native passband through phase-stepped acquisition and demodulation across shifted passbands (discrete filling). Note, the lateral and axial resolution improvements come at the cost of multi-angle/multi-pitch acquisitions and require interferometric stability.

In this paper, we report on a 5-illumination component implementation of c-SIM which provides enhanced optical sectioning compared to conventional quadrupolar illumination. This technique takes advantage of an on-axis (zero-order) illumination beam that introduces additional interference terms into the excitation fields, producing stronger axial modulation compared with the 4-beam case. This provides enhanced rejection of out-of-focus light in addition to a 2-fold lateral resolution improvement compared to the optical system’s original passband while also avoiding tomographic angle scanning or pitch modulation. Since images are recorded as intensities due to coherently scattered light, our approach avoids the complexities of interferometric field detection. Since 5-beam c-SIM technique achieves enhanced optical sectioning by suppressing out-of-focus information detected through the optical system’s original axial passband, it is generally applicable to 2D imaging scenarios that require fast acquisition rates since the number of frames required for reconstruction is reduced by an order of magnitude compared to label-free 3D volumetric approaches that detect coherent signals.

## 2. Background and Methods

Our approach to c-SIM relies on coherent excitation with 2D structured illumination (SI) patterns which modulate sample frequency information by the orthogonal spatial frequency vectors *ω*_0,*x*_, *ω*_0,*y*_. The SI intensity patterns may be expressed as:

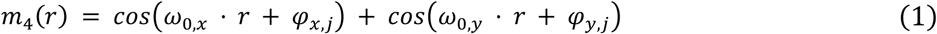

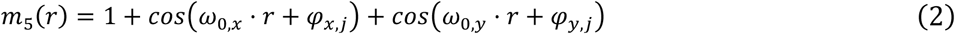

for the 4- and 5-beam cases, respectively. Here, *ω*_0_ = 2*π/N*where *N*is the period of the SLM grating in pixels. Note that the DC term in (2) includes self-interference, |E_0_|^2^, of the on-axis (zero-order) field. Analogous to fluorescence SIM, the phases, (*φ*_*x,j*_, *φ*_*y,j*_), dictate the position of the *j*^*th*^ periodic SI pattern and sets of phase-shifted images acquired at predetermined phases may be described as a linear system. The raw image spectra contain multiplexed, shifted copies of the sample spatial frequencies which can be separated into constituent components through harmonic demodulation (described in more detail below). Representing the illumination field spectrum as a sum of discrete plane-wave components clarifies that the phase factors act as complex weights for the spatial frequency vectors at ±*ω*_0,*x*_, ±*ω*_0,*y*_ for the 4-beam case and *ω* = 0, ±*ω*_0,*x*_, ±*ω*_0,*y*_ for 5-beam illumination.

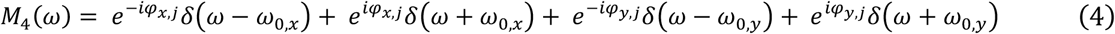

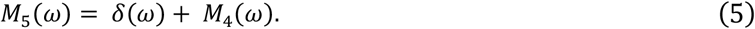

In (4) and (5), the 2*π* normalization factors have been omitted. The number of phase-stepped acquisitions required in each experiment, and thus, the dimensionality of the linear systems, is governed by the coherent spatial frequency distribution recorded at the detector given by

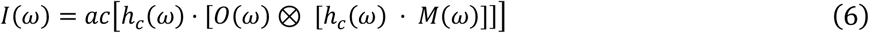

where *h*_*c*_(*ω*) is the coherent transfer function (CTF) defined by a circular pupil extending out to the frequency cutoff, *ω*_*max*_, governed by the excitation objective NA. Here, *O*(*ω*) and *M*(*ω*) represent the object and SI spatial frequency distributions, respectively, and we assume detection is through the excitation optic. In (6), *a* and ⊗ are the autocorrelation and convolution operations, respectively. Substituting (4) and (5) into (6), expanding, and grouping similar phase terms (see Supplemental Information) yields expressions for the spatial frequency distributions for a given phase pair, (*φ*_*x,j*_, *φ*_*y,j*_), in the 4- and 5-beam c-SIM experiments. The unique spectral terms generated due to the spatial phase modulation for the more general case of 5-beam illumination are summarized in Table S1. The illumination results in 13 unique terms requiring the same number of SI image acquisitions to describe the enhanced resolution components completely. In the case of 4-beam illumination, terms 6-9 (Table S1) are absent since they are zeroth order - ±x or ±y cross terms; only 9 phase-stepped acquisitions are required for 4-beam illumination.

The experimental setup for carrying out 4- and 5-beam c-SIM is described in Fig. 1a. The laser source is derived from the fundamental output of a femtosecond ti:sapphire oscillator (Griffin, KM Labs) operating at a carrier wavelength of 800 nm (40 nm bandwidth, ∼ 30 fs) which passes through a 50:50 beamsplitter and is then reflected from a spatial light modulator (EXULUS-HD3HP, Thorlabs, Inc.). We utilize a pulsed laser in anticipation of nonlinear implementations of c-SIM. Note, the source beam is pre-conditioned for spectral dispersion using a pair of chirped mirrors and spatially filtered using a pinhole. The SI patterns are formed by uploading binary 2D square lattice gratings onto the SLM. The SLM plane is relayed by a 30 cm–10 cm two-lens system (≈1/3x magnification) to an intermediate plane, and a 20 cm tube lens images this relay onto the objective back focal plane. Demagnification pre-scales the angular separation of the diffracted orders such that it matches the objective pupil diameter after the tube lens. As depicted in Fig. 1b and c, a mask placed at the intermediate plane is used to isolate the ±1^st^ diffraction orders and optionally the zero-order beam emanating from the SLM. The zero-order beam is toggled off/on for 4/5 beam c-SIM measurements allowing a direct comparison of these configurations. Images are acquired by a CMOS camera (LP126MU, Thorlabs Inc.) in the reflected geometry using a beamsplitter placed in the infinity space.

**Figure 1.**
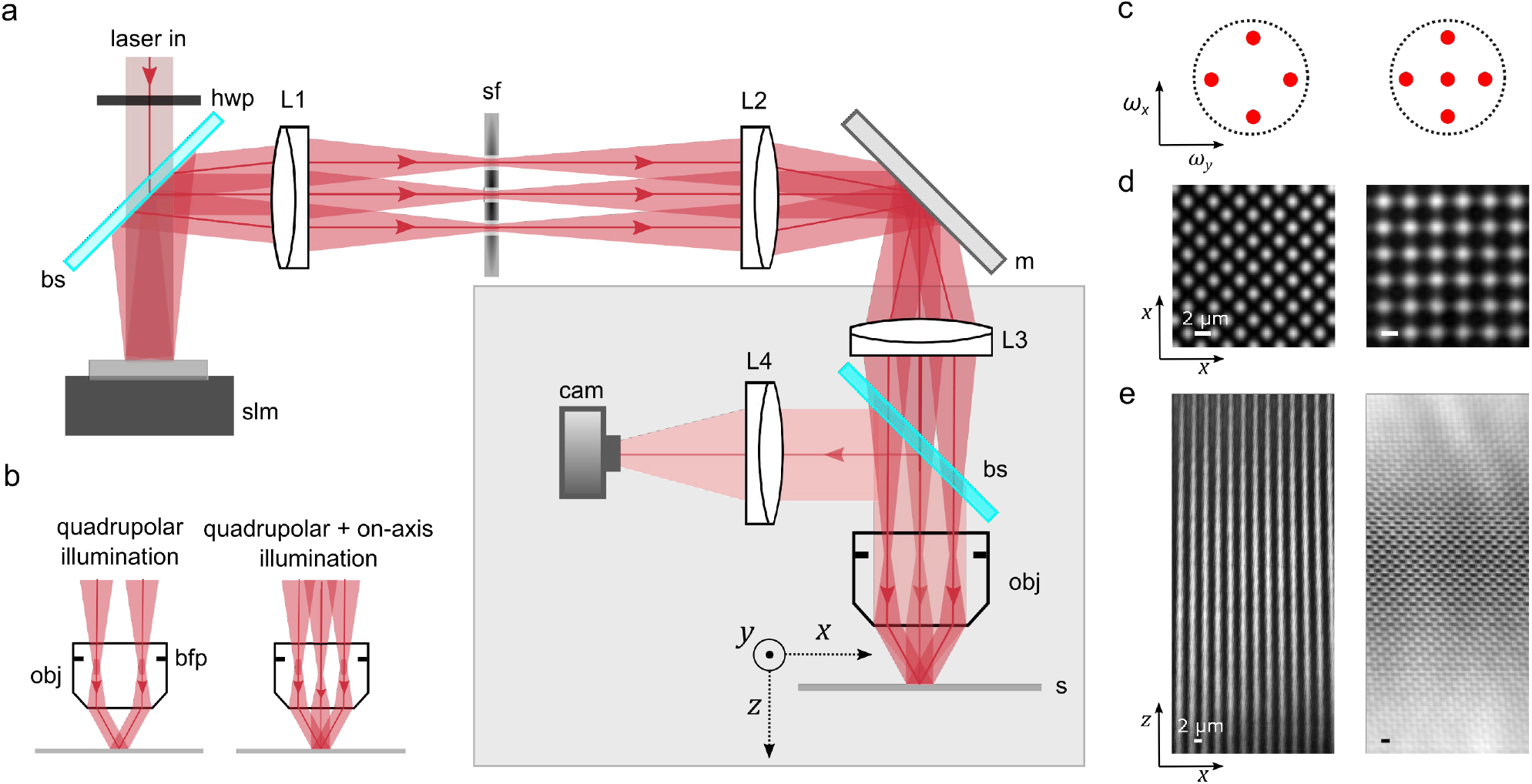
Experimental setup for coherent structured illumination microscopy (c-SIM). In (a), hwp: half-wave plate; bs: beamsplitter; slm: spatial light modulator; L1, L2, L3, L4: lenses (f = 30, 10, 20, 18 cm); sf: spatial filter; m: mirror; obj: objective lens; s: sample; cam: CMOS camera. b) 4- and 5-beam c-SIM configurations for wide-field microscopy, c) corresponding schematics of the back focal planes, and d) real-space illumination patterns at one phase point in the *x* − *y* plane (sample plane). e) Profiles of the 4-(left) and 5-beam (right) SI patterns acquired along the *x* − *z* plane. The *z* range is 600 µm.

The demodulation procedure is based on illumination with sets of SI patterns with known phases. Phase sets are chosen from an *N*× *N*grid composed of discrete steps:

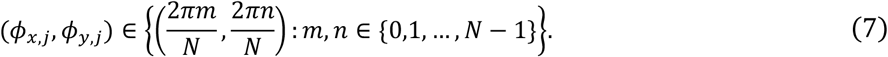

Letting *y*(*ω*) = [*Y*_1_(*ω*), … , *Y*_*j*_(*ω*)]^*T*^ be the measured spectra at phase pairs *Φ*_*x,j*_, *Φ*_*y,j*_, and *c*(*ω*) be the unknown harmonic components, then *Φ* is chosen such that

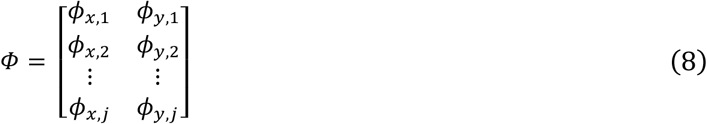

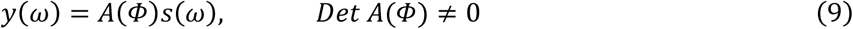

is satisfied. In practice, the SLM pixel pitch is chosen to correspond to *N*which ensures the phase steps are integer multiples of the pitch. Since the angular separation of the diffracted orders also depends on *N*, it is adjusted to maximize *ω*_0_ for different optics.

Images of the 4- and 5-beam illumination structure at one phase point recorded on a flat mirror are shown in Fig. 1d. The superposition of square and diagonal frequencies in the 4-beam case gives rise to the triangle-like pattern, although both cases have the same carrier frequency and four-fold symmetry. This is also clear from phase maps derived from the raw image spatial spectra (see Fig. S1). Here, the phase, *Ar A*[*Y*_*i*_(*ω*)*/ Y*_*j*_(*ω*)], is plotted for several *i, j* combinations showing the evolution of terms for isolated channels or their mixtures. The 5-beam SI pattern exhibits additional axial structure as shown in Fig. 1e.

Our reconstruction procedure follows the basic steps given in Ref. 6. Raw SI images are converted into image spectra and separated into constituent components using *c*(*ω*) = *A*^−1^ *y*(*ω*) and the programmed phases. Cross-correlation of the unmixed terms with the zero-order spectrum is used to determine *ω*_0,*x*_, *ω*_0,*y*_. The images are padded by a factor-of-2 in the frequency domain, back-transformed into position space and multiplied by real-space phase gradients corresponding to the spatial phase modulation in pixel space determined in the previous step. The final reconstruction is carried out by summing the modulated real-space images after weighting the spectra with complex factors.

## 3. Experimental Results

### 3.1 Sub-diffraction imaging with 5-beam c-SIM

To demonstrate the lateral resolution improvement attainable with c-SIM, experimental images acquired through on-axis illumination with a single beam and through 5-beam c-SIM are compared in Figure 2a. These experiments utilize a modest 4× objective (NA 0.16) to avoid generating transverse polarization components. The SLM pitch of (*N*= 5) pixels and corresponding spatial frequency vectors are close to *ω*_*max*_. A visual comparison of the WF image and c-SIM reconstruction (Fig. 2a) of a silicon atomic force microscope target shows a qualitative improvement in the sharpness of the individual elements and a noticeable suppression of the diffractive ringing surrounding them. Discrete Fourier transforms (DFT) of the images are shown in Fig. 2b. These data are superimposed with CTF masks synthesized using the central illumination wavelength (800 nm) and an on-axis CTF simulated using a scalar Fraunhofer diffraction model. To visualize the extended c-SIM aperture, the on-axis CTF was translated according to the shift vectors established using the cross-correlation procedure described above.

**Figure 2.**
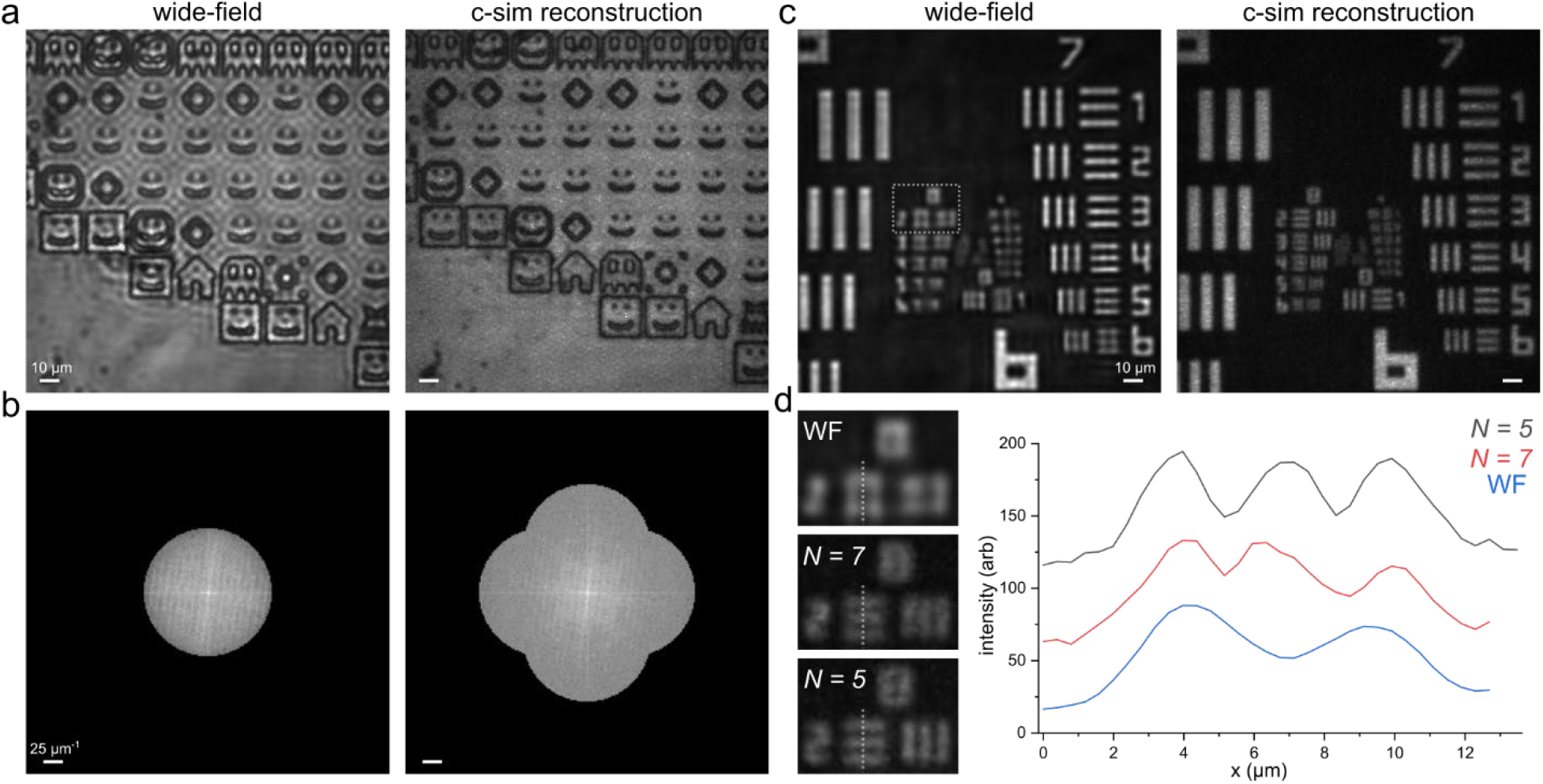
a) Experimental data comparing wide-field (left) and c-SIM images (right) of a coherently scattering sample. b) Spatial spectra (magnitudes) corresponding to the images in (a). The data are superimposed with CTFs simulated using the parameters extracted from the experimental reconstruction to highlight the extended frequency support for c-SIM (right). Note, the c-SIM experiment utilized *NN*= 5 SLM pixels for the period of the spatial frequency modulation which is close to the frequency cutoff for the objective lens. c) Analysis of the lateral resolution improvement using wide-field and c-SIM images acquired on a test resolution target. d) Influence of SLM pitch on the attainable resolution (left) and line profiles of group 8 element 2 (right).

To quantify the lateral resolution improvement, images of a USAF test resolution target were recorded using WF and 5-beam c-SIM (Fig. 2c). Using Michelson contrast modified for coherent detection[16] as a metric for the frequency cutoff yields a 1.7-fold increase in lateral resolution for c-SIM (*N*= 5) compared to on-axis illumination (see Fig. S2). To visualize the attainable resolution, we provide a comparison of c-SIM images recorded on group 8 element 2 for different SLM pitches (*N*= 5, 7) in Fig. 2d. Plots of line profiles acquired on the horizontal bar set show the increased intensity modulation as the period of the SLM grating decreases, and thus, the modulation spatial frequency correspondingly increases. Since the extended aperture and test resolution target are co-aligned along the horizontal and vertical axes in spatial frequency space, the resolution improvement in *x* and *y* is similar.

### 3.2 Optical sectioning in 4- and 5-beam c-SIM

The 4 and 5-beam c-SIM implementations may be compared by toggling the zero-order beam on or off while keeping other parameters fixed. Although less patterned acquisitions are required to reconstruct the quadrupolar data, 9 pairs from the set of 13 phases chosen from (7) may be found such that *DeD A*(*Φ*) ≠ 0. As such, the same SLM gratings are used in both experiments. In Fig. 3, we present on-axis, 4-beam, and 5-beam c-SIM images of 2 µm latex beads dispersed on a silver-coated coverslip. Comparison of the c-SIM images to the wide-field reveals a similar factor of lateral resolution improvement as demonstrated in Fig. 2. The principle difference is enhanced suppression of the diffractive ringing in the 5-beam experiment compared to the 4-beam case as evidenced by the individual sphere profiles. Since 2 µm is below the on-axis diffraction limit, the single sphere profiles in the wide-field image exhibit lateral broadening and out-of-focus diffractive effects. The coherent defocus artifacts are somewhat less apparent in the 4-beam data while the 5-beam images more clearly resolve the individual sphere profiles and recover the FWHM of 2 µm in the lateral dimension. To analyze this further, axial slices were acquired by scanning the objective lens through the focal plane of the beads while carrying out 4 and 5-beam c-SIM reconstructions at each axial slice. Full *x* − *z* sections acquired at the *x* position indicated by the red line on the wide-field image are shown in Fig. 3b along with line cuts centered on the single sphere (Fig. 3c). While the c-SIM profiles appear intensity-like, the wide-field profile exhibits familiar through-focus coherent interference behavior which produces oscillatory lobes that manifest as light and dark fringes. The wide-field, 4-beam, and 5-beam axial profiles exhibit progressively stronger suppression of the defocus lobes that appear on either side of the central bead peak. Gaussian fits of the c-SIM curves yield an effective 40% axial sharpening of the bead profile for the 5-beam case.

**Figure 3.**
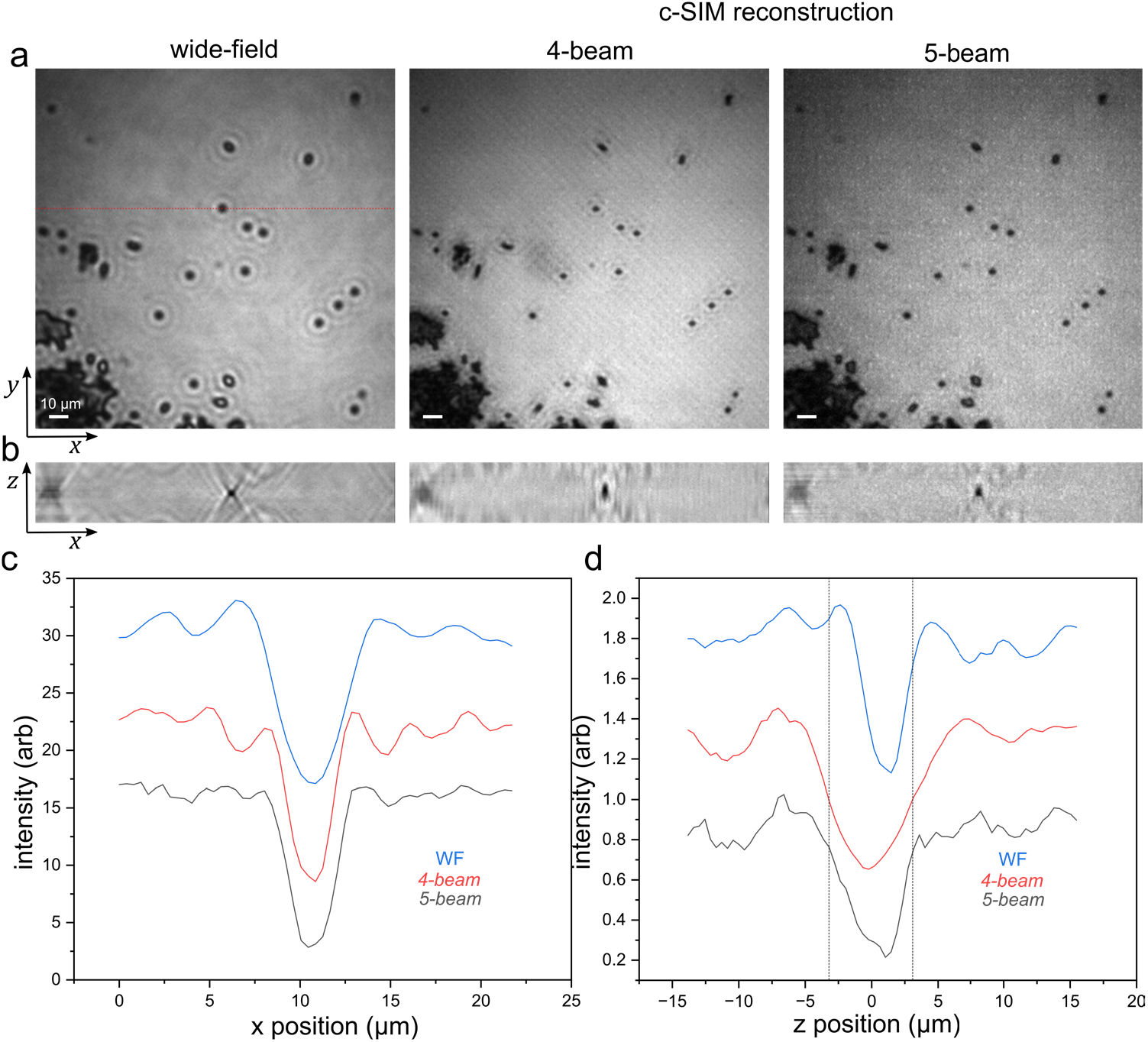
Comparison of 4- and 5-beam c-SIM. a) On-axis and c-SIM reconstructions of 2 µm latex beads using a 4× (NA 0.16) objective. b) Axially resolved 2D wide-field and c-SIM reconstructions recorded at the *y* plane indicated by the red line in (a), c-d) Lateral and axial profiles of the central bead appearing as indicated in (a). In (d), the droplines correspond to the FWHM of the 4-beam profile.

As mentioned above, the coherently detected wide-field modality provides zero axial gating since the on-axis CTF is infinitesimally narrow in 3D spatial frequency space. A limited optical sectioning effect is observed in the 4-beam experiment since the interference region is finite and dictated by the beam obliquity. Visualizing the 4- and 5-beam SI profiles in the *x* − *z* plane (Fig. 1e) shows the stronger axial modulation that appears due to the presence of additional interference components in the illumination (see Table S1). Note, the axial sections presented are uncoupled stacks; 2D c-SIM is carried out at different *z* - planes to analyze the relative optical sectioning strength. The additional zero-order term in the 5-beam experiment provides axial modulation that could be included in the reconstruction to obtain a true 3D advantage as in 3-beam fluorescence SIM,[6] but the pitch would also need to be scanned to fully capture the other spatial frequencies.[20]

### 3.3 c-SIM of live cells

Fig. 4 summarizes live-cell c-SIM experiments carried out on *Vacuoliviride crystalliferum* microalgae.[26] In this system, vegetative cells are characterized by a large central vacuole, refractile crystalline granules, and parietal chloroplasts. The distribution of absorption and scattering interior cell features, presence of phase-only components, and the cell thickness make it an ideal specimen for demonstrating the utility of the 4- and 5-beam c-SIM modalities. In these experiments, the cells are suspended in low melt temperature agar to prevent diffusion, drop deposited on a microscope slide, and sealed with cover glass. The wide-field image is marked by broadened internal cell features and coherent defocus ringing around the cells, similar to the on-axis images of latex spheres (Fig. 3a). Defocus effects are especially noticeable in the upper region of the image where the rings from neighboring cells coalesce, obscuring the cell boundaries. In the c-SIM images, the cell boundaries are qualitatively sharper, and the stronger optical sectioning strength of the 5-beam configuration is operative as the regions between cells are much less obscured by diffractive ringing. Although the background in the 5-beam image appears to exhibit residual patterning, the intracellular features appear more distinctive due to axial gating in the same manner as the cell boundaries.

**Figure 4.**
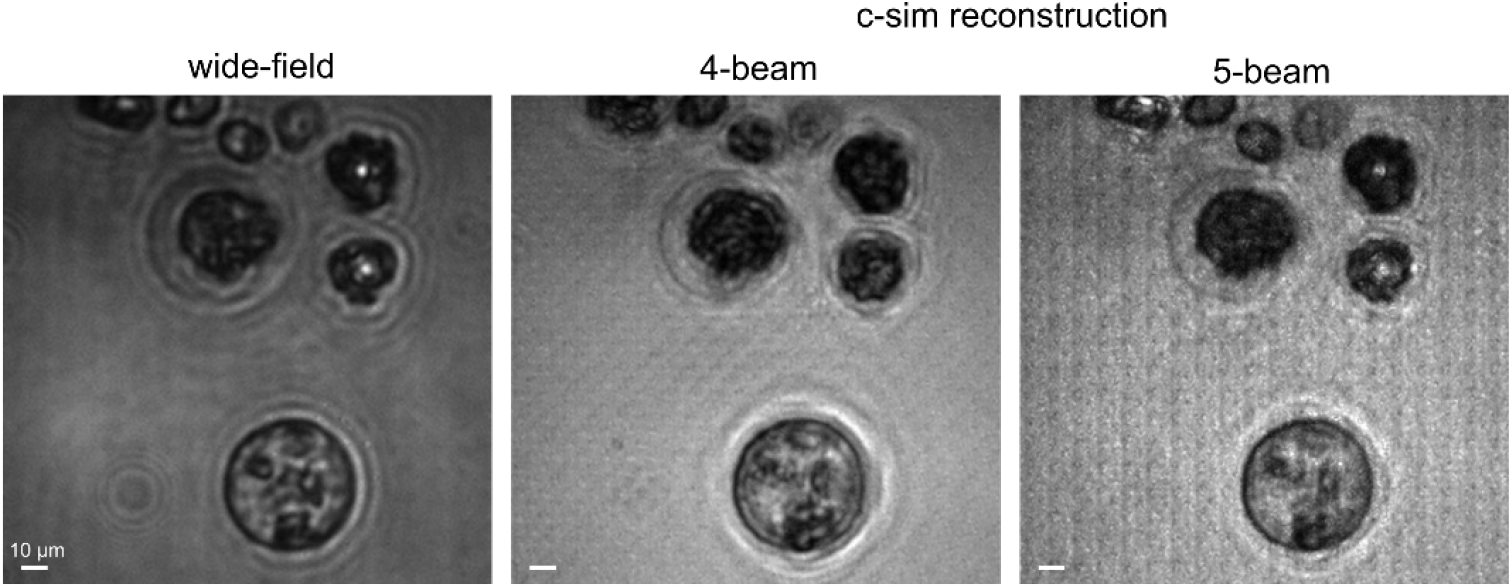
c-SIM images acquired on live *Vacuoliviride crystalliferum* microalgae under 10× (NA 0.4) illumination.

### 3.4 Implications of femtosecond illumination

There are several practical considerations that the utilization of an ultrafast pump introduces. First, the pump bandwidth has the effect of CTF averaging since *ω*_*max*_ ∝ 1*/*λ. As a result, the intensity modulation frequency dependence (see Fig. S2) is sloped. Moreover, the large bandwidth causes the modulation peaks in the raw image spatial spectra to become smeared, effectively reducing modulation contrast at the highest frequencies. Second, the illumination beams pick up angular dispersion after the SLM. Since the illumination beams are spectrally dispersed at the back focal plane, large bandwidth pulses will exhibit elongated spatial profiles which limits the maximum spatial frequency modulation attainable without clipping. This amounts to a pulse duration – spatial resolution tradeoff in the case where the angular dispersion is not compensated for.

## 4. Discussion

A comparison of 4- and 5-beam c-SIM results on the same samples highlight key differences between these modalities. Since the spatial modulation carrier frequency is the same, the extended aperture and, consequently, the lateral resolution cutoff is equivalent in both configurations. Our basic reconstruction framework utilizes a complex scalar weight factor per demodulation channel. As such, compared to 4-beam c-SIM, the 5-beam implementation tends to overestimate lower frequencies. This can be addressed by utilizing spatial frequency dependent weights although this increases the complexity of the parameter determination and image reconstruction.[6, 8]

Qualitative differences in 4- and 5-beam c-SIM images may be rationalized by considering optical sectioning effects. While on-axis illumination provides no optical sectioning under coherent illumination and detection conditions, 4-beam c-SIM shows suppression of out-of-focus diffractive ringing since the interference between oblique beams is confined to their region of overlap. Since 5-beam c-SIM adds additional, axially dependent, interference terms into the illumination (Fig. 1e), a comparatively strong optical sectioning effect is operative as evidenced by suppression of the oscillatory features in the bead lateral profiles (Fig. 3c), and the narrower effective bead axial profile (Fig. 3d). Note, in the experiments presented, the axial bandwidth is the same in the 4- and 5-beam configurations.

## 5. Conclusion

We have presented complementary 4- and 5-beam frameworks that provide 2-fold lateral resolution improvements in wide-field coherent structured illumination microscopy. Here, 2D sample reflectance images are recorded as intensities, simplifying the optical setup and reconstruction procedure compared to complex field detection. We have shown that the addition of an on-axis beam in c-SIM produces comparatively enhanced optical sectioning which is accounted for by considering the additional spectral terms that arise due to zero-order /oblique beam interference. Measurements carried out on microalgae demonstrate the utility of c-SIM for suppression of coherent defocus effects. Our intensity-based imaging approach provides enhanced optical sectioning and qualitative amplitude images but does not directly access phase information. Various phase retrieval techniques could be implemented such as those based on the transport-of-intensity equations.[27, 28] This approach would recover phase-and-amplitude information using the experimental setup without modification. Note, although the signals presented here are linear in pump fluence, detecting the coherent scattering channel utilizing a femtosecond pump source paves the way for nonlinear implementations of c-SIM.

## Acknowledgements

The authors would like to thank Jerry Kuper (University of Rochester) for helpful discussions and contributions to laboratory operations during this work. The authors kindly thank Jana Pilátová (Charles University) for providing the microalgae.

## Data Availability

The data that support the findings of this study are available upon reasonable request.

## Supplemental Documentation

See Supplement 1 for supporting content.

