## Supplement 1 for "Coherent Structured Illumination Microscopy with Enhanced Optical Sectioning"

The structured illumination components for the 5-beam case may be found by substituting equations (4) and (5) into (6), expanding, and grouping similar phase terms. This yields an expression for the coherent spatial frequency distribution recorded at the detector as given below in Eqn. S1. Here, $ac$ and $*$ designate auto-correlation and cross-correlation, respectively. The spectral phase terms are summarized in Table S1. Note, the 1st harmonic $x$ and $y$ components (terms 6-9 in Table S1) are absent in 4-beam illumination.

$$
\begin{aligned}
I(\omega) = &\begin{bmatrix} ac\left[h_c(\omega)\, M\left(\omega-\omega_{0,x}\right)\right] + ac\left[h_c(\omega)\, M\left(\omega+\omega_{0,x}\right)\right] + \\ ac\left[h_c(\omega)\, M\left(\omega-\omega_{0,y}\right)\right] + ac\left[h_c(\omega)\, M\left(\omega+\omega_{0,y}\right)\right] \end{bmatrix} + \\
&e^{-i2\phi_x}\left[h_c(\omega)\, M\left(\omega-\omega_{0,x}\right) * h_c(\omega)\, M\left(\omega+\omega_{0,x}\right)\right] + \\
&e^{i2\phi_x}\left[h_c(\omega)\, M\left(\omega+\omega_{0,x}\right) * h_c(\omega)\, M\left(\omega-\omega_{0,x}\right)\right] + \\
&e^{-i2\phi_y}\left[h_c(\omega)\, M\left(\omega-\omega_{0,y}\right) * h_c(\omega)\, M\left(\omega+\omega_{0,y}\right)\right] + \\
&e^{i2\phi_y}\left[h_c(\omega)\, M\left(\omega+\omega_{0,y}\right) * h_c(\omega)\, M\left(\omega-\omega_{0,y}\right)\right] + \\
&e^{-i\phi_x}\begin{bmatrix} h_c(\omega)\, M\left(\omega-\omega_{0,x}\right) * h_c(\omega)\, M(\omega) \\ h_c(\omega)\, M(\omega) * h_c(\omega)\, M\left(\omega+\omega_{0,x}\right) \end{bmatrix} + \\
&e^{i\phi_x}\begin{bmatrix} h_c(\omega)\, M\left(\omega+\omega_{0,x}\right) * h_c(\omega)\, M(\omega) \\ h_c(\omega)\, M(\omega) * h_c(\omega)\, M\left(\omega-\omega_{0,x}\right) \end{bmatrix} + \\
&e^{-i\phi_y}\begin{bmatrix} h_c(\omega)\, M\left(\omega-\omega_{0,y}\right) * h_c(\omega)\, M(\omega) \\ h_c(\omega)\, M(\omega) * h_c(\omega)\, M\left(\omega+\omega_{0,y}\right) \end{bmatrix} + \\
&e^{i\phi_y}\begin{bmatrix} h_c(\omega)\, M\left(\omega+\omega_{0,x}\right) * h_c(\omega)\, M(\omega) \\ h_c(\omega)\, M(\omega) * h_c(\omega)\, M\left(\omega-\omega_{0,x}\right) \end{bmatrix} +
\end{aligned}
\tag{S1}
$$

$$e^{i\left(\phi_x-\phi_y\right)}\begin{bmatrix}h_c(\omega)\,M\left(\omega-\omega_{0,x}\right)*\;h_c(\omega)\,M\left(\omega-\omega_{0,y}\right)\\h_c(\omega)\,M\left(\omega+\omega_{0,y}\right)*\;h_c(\omega)\,M\left(\omega+\omega_{0,x}\right)\end{bmatrix}+$$

$$e^{-i\left(\phi_x-\phi_y\right)}\begin{bmatrix}h_c(\omega)\,M\left(\omega-\omega_{0,x}\right)*\;h_c(\omega)\,M\left(\omega+\omega_{0,y}\right)\\h_c(\omega)\,M\left(\omega-\omega_{0,y}\right)*\;h_c(\omega)\,M\left(\omega+\omega_{0,x}\right)\end{bmatrix}+$$

$$e^{i\left(\phi_x+\phi_y\right)}\begin{bmatrix}h_c(\omega)\,M\left(\omega+\omega_{0,x}\right)*\;h_c(\omega)\,M\left(\omega-\omega_{0,y}\right)\\h_c(\omega)\,M\left(\omega+\omega_{0,y}\right)*\;h_c(\omega)\,M\left(\omega-\omega_{0,x}\right)\end{bmatrix}+$$

$$e^{-i\left(\phi_x+\phi_y\right)}\begin{bmatrix}h_c(\omega)\,M\left(\omega-\omega_{0,y}\right)*\;h_c(\omega)\,M\left(\omega-\omega_{0,x}\right)\\h_c(\omega)\,M\left(\omega+\omega_{0,x}\right)*\;h_c(\omega)\,M\left(\omega-\omega_{0,y}\right)\end{bmatrix}$$

| *term* | *component* | *phase* | *origin* |
|---|---|---|---|
| 1 | DC | 1 | DC / 0th order |
| 2–3 | ±2x | $e^{\pm i2\phi_x}$ | 2nd harmonic (x) |
| 4–5 | ±2y | $e^{\pm i2\phi_y}$ | 2nd harmonic (y) |
| 6–7 | ±1x | $e^{\pm i\phi_x}$ | fundamental / 1st harmonic (x) |
| 8–9 | ±1y | $e^{\pm i\phi_y}$ | fundamental / 1st harmonic (y) |
| 10–11 | ±(x−y) | $e^{\pm i(\phi_x - \phi_y)}$ | diagonal mixed harmonic (difference) |
| 12–13 | ±(x+y) | $e^{\pm i(\phi_x + \phi_y)}$ | diagonal mixed harmonic (sum) |

Table S1. Spectral terms arising from 2D spatial phase modulation under 5-beam illumination.

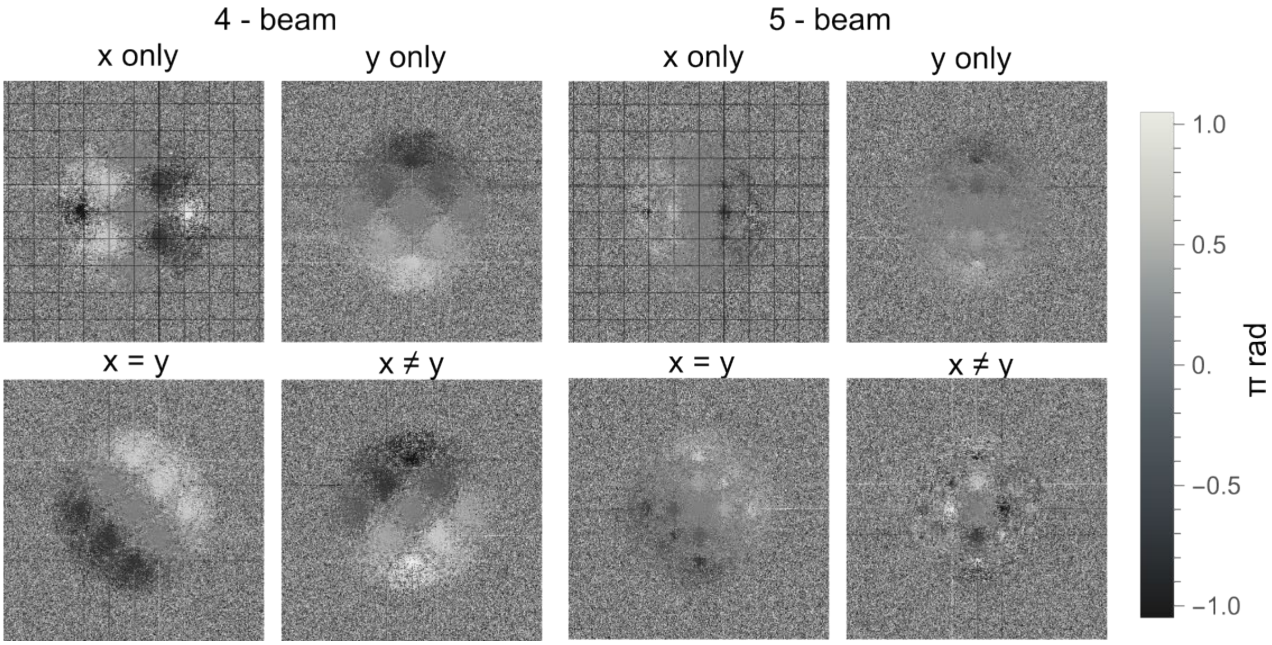


Figure S1. Phases maps for several structured illumination components for each c-SIM configuration showing the association between real-space pattern translations and the amplitude of various frequency components for $x, y, x = y, x \neq y$. The grid is spaced at modulation carrier frequency $\omega_0$.

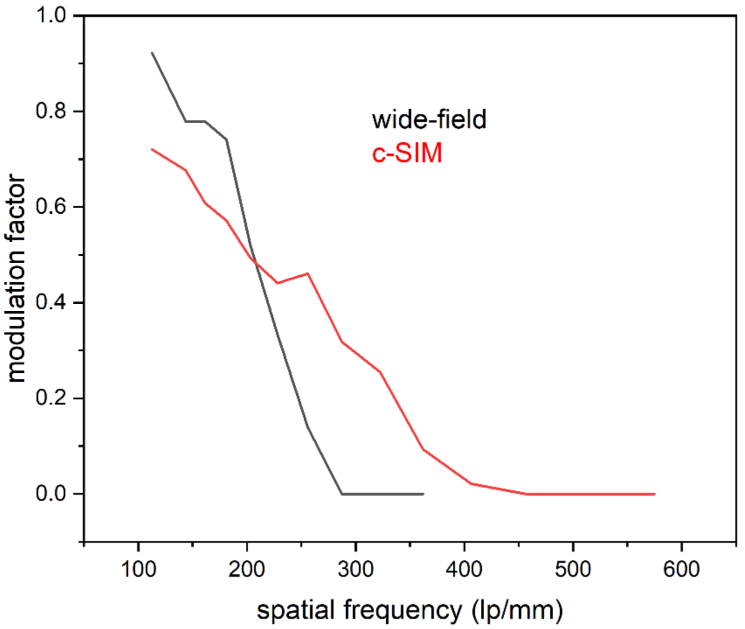


Figure S2. Intensity modulation recorded as a function of frequency for WF and c-SIM ($N = 5$) under 4× (NA 0.16) illumination. Data extracted from the images presented in Fig. 2c.
